# Predicting the impact of non-coding variants on DNA methylation

**DOI:** 10.1101/073809

**Authors:** Haoyang Zeng, David K. Gifford

## Abstract

DNA methylation plays a crucial role in the establishment of tissue-specific gene expression and the regulation of key biological processes. However, our present inability to predict the effect of genome sequence variation on DNA methylation precludes a comprehensive assessment of the consequences of non-coding variation. We introduce CpGenie, a sequence-based framework that learns a regulatory code of DNA methylation using a deep convolutional neural network and uses this network to predict the impact of sequence variation on proximal CpG site DNA methylation. CpGenie produces allele-specific DNA methylation prediction with single-nucleotide sensitivity that enables accurate prediction of methylation quantitative trait loci (meQTL). We demonstrate that CpGenie prioritizes validated GWAS SNPs, and contributes to the prediction of functional non-coding variants, including expression quantitative trait loci (eQTL) and disease-associated mutations. CpGenie is publicly available to assist in identifying and interpreting regulatory non-coding variants.

## 1 Introduction

A significant portion of the disease and trait-associated variants revealed by genome-wide association studies (GWAS) reside in the non-coding genome where they alter cellular activities and organism phenotype by changing gene regulation [1–3]. While GWAS studies can identify thousands of loci that are associated with traits, they are typically underpowered to identify the exact causal variants for a trait of interest, and further analysis of the potential functional consequence of each variant must be performed. Computational methods that analyze candidate variants for their potential contribution to a phenotype of interest are known as variant prioritization methods. Variant prioritization methods that accurately predict which variants influence proximal regulatory elements and thus gene regulation are valuable tools.

Previous variant prioritization methods have considered a diverse set of functional signals [4–8], including DNase hyper-sensitivity sites (DHS), histone modifications, and transcription factor binding. However, DNA methylation, an important epigenetic state that is involved in the regulation of key biological processes [9–14] and encodes cellular state information not contained in other epigenetic marks [15, 16] has been largely overlooked. In the few methods where it is considered [5], DNA methylation is used as a low-resolution regional feature that is not allele-specific. While sequence-based methods for DNA methylation exist [17–20], there is no published method to predict the impact of sequence variants on methylation, which makes it difficult to incorporate DNA methylation in functional variant prioritization.

We introduce CpGenie (Figure 1), a deep-learning model that (i) learns a regulatory code of DNA methylation, (ii) predicts the methylation status of a CpG site from the flanking sequence at a single-nucleotide sensitivity and (iii) produces high-confidence predictions of non-coding variants that module DNA methylation. We find that CpGenie predicts the impact of sequence variants on DNA methylation with an accuracy that surpasses existing methods for functional variants prioritization. CpGenie also identifies the direction of impact of meQTLs that result in an allelic imbalance of DNA methylation, and prioritizes meQTLs over variants that exhibit no effect on DNA methylation with accuracy higher than alternative methods. We show that predictions from CpGenie improves the prediction of expression quantitative trait loci (eQTLs) and disease-associated variants by providing functional information complementary to other data type. In addition, we find that the sequence determinants learned by CpGenie correspond to the binding motifs of proteins known for their involvement in the regulation of DNA methylation state. We provide CpGenie as open source software available at http://cpgenie.csail.mit.edu.

## 2 Materials and Methods

### 2.1 CpGenie implementation

#### Me-CpG-prediction

We implemented a three-layer convolutional neural network with Dropout and max-norm regularization. Our implementation utilizes the Keras library (https://keras.io). To cope with the differences in sample sizes and protocol, we used slightly different network parameters for models trained on RRBS data from ENCODE and the pool-based bisulfite sequencing data from [21]. Detailed network structure can be found in Supplemental Table 5. Hyper-parameters, such as learning rate and Dropout ratio, are tuned in a standard cross-validation fashion with test set completely held out. As input, each DNA sequence of length *L* is converted into a 2-D matrix of size 4 × *L*, where each column is a one-hot vector encoding the presence of the four DNA nucleotides A, C, G, and T.

#### Variant-prediction

Given a sequence variant, we predict the methylation status of all CpGs within 500 bp with either of the variant allele. The maximum, mean and sum of the methylation level of adjacent CpGs are reported for each allele. In case where no CpG resides in the 500bp vicinity of the given variant, a pseudo-methylation level of 0.001 is reported. We evaluate the impact of the variant by calculating the change

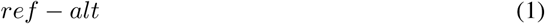

in the sum/mean/max methylation level, and the change of log odds

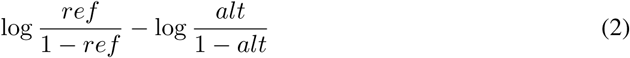

in the mean/max methylation level of nearby CpG sites, resulting in five features for each variant.

### 2.2 High-throughput DNA methylation data

The 50 RRBS datasets of immortal cell lines, including GM12878, were downloaded from ENCODE website (https://www.encodeproject.org/). We merged multiple replicates for the same experiments, and where a CpG exists in all replicates we merged the counts of methylated and unmethylated reads and re-calculated the percentage of methylation. We further applied a minimum-read cutoff of 10 to filter out unreliable samples. Samples from chromosome 1-9 and chromosome 14-22 were used for training, samples from chromosome 12-13 were used for hyper-parameter tuning and model selection, and the rest of the data were held-out for testing.

The raw allele-specific DNA methylation data were obtained from Fraser lab (personal communication). They surveyed 823,726 SNP-CpG pairs, among which 2,379 are meQTLs. After filtering out CpGs with read counts less than 10, the whole dataset was split into training, validation and testing set in the same way as previously described for RRBS data. Test set was completely held out from training. For simplicity, only methylation levels corresponding to the reference allele were used in the training and evaluation of Me-CpG-prediction. In the analysis of variant-prediction, only meQTL and allele-specific methylation data from the held-out chromosome 10 and 11 were used.

### 2.3 Methylation prediction comparison with random forest

We counted the frequency of each possible 4-mer in the 1001bp sequence centered at a CpG with JELLYFISH [22] (version 2.2.6, https://github.com/gmarcais/Jellyfish/releases), generating 256 features for each sample. We used the random forest implementation in sciki-learn Python package (http://scikit-learn.org/stable/modules/generated/sklearn.ensemble.RandomForestClassifier.html).

### 2.4 meQTL prioritization comparison with existing methods

We downloaded DeepSEA (ver. 0.93) from http://deepsea.princeton.edu/, GWAVA (ver. 1.0) from http://www.sanger.ac.uk/resources/software/gwava/, deltaSVM from http://www.beerlab.org/deltasvm/, and Basset from https://github.com/davek44/Basset. For CADD (ver. 1.3), we used the online webserver (http://cadd.gs.washington.edu/).

For deltaSVM and Basset, the predictions of which are cell-line specific and direction-included, we used the absolute value of the predictions for the same type of cell line (LCL, lymphoblastoid cell line) from which the meQTL were discovered from. For deltaSVM, we used the gkmSVM weights trained on GM12878 DNase Hyper-sensitive Sites (DHS). For Basset, we used the absolute SAD (SNP Accessibility Difference) scores predicted for GM12878.

### 2.5 Functional variant prioritization

The variants in strong linkage disequilibrium with rs1427407, rs12740374, rs10737680, rs7705033 are determined by finding all variants with *r*^2^ = 1 in HaploReg (http://archive.broadinstitute.org/mammals/haploreg/haploreg.php, v4.1). In the case of rs1427407 where no variants match the criteria, we find all variants with *r*^2^ >= 0.8 with it.

We obtained the eQTL and GWAS SNPs datasets, as well as their corresponding five negative sets from the supplementary tables in Zhou et al. [6]. Four of the five negative sets were constructed by finding, for each positive variant, the closest SNP in the full set, 20%, 4% and 0.8% random subset of 1000 Genome variants with minor allele frequency distribution matched to the positive set. The mean distance to the positive set is 360 bp, 1,400 bp, 6,300 bp and 31k bp for these four negative sets respectively. The fifth negative set was constructed by sampling 1,000,000 non-coding 1000 Genome SNPs with minor allele frequency distribution matched to the positive set.

For each variant in the positive and negative set, we applied variant-prediction to generate DNA methylation features that describe the impact on the proximal DNA methylation levels in the 50 ENCODE RRBS dataset, resulting in 250 features for each variant. We further kept only the absolute value of each feature. As described in Zhou et al., for each negative set we trained *L*_2_-regularized logistic regression models on CpGenie and DeepSEA features respectively using scikit-learn library (http://scikit-learn.org/stable/modules/generated/sklearn.linear_model.LogisticRegressionCV.html). The performance was evaluated with 10-fold cross-validation. For CADD, GWAVA and Funseq2, the auROC reported in Zhou et al. [6] was directly used as we tested on the same dataset.

To interpret the feature importance, we trained a random forest classifier on the same tasks as above with all the features normalized to have mean 0 and variance 1 before training. We used random forest implementation in scikit-learn library (http://scikit-learn.org/stable/modules/generated/sklearn.ensemble.RandomForestClassifier.html) in which the feature importance is calculated as the “mean decrease impurity” defined as total decrease in node impurity averaged over all trees of the ensemble [23]. To interpret the features with markedly higher importance, for each model we plotted the sorted feature importance and identified an importance-cutoff corresponding to the ‘elbow-point’ in the importance distribution. The top features in each model are defined as the ones with importance higher than the corresponding cutoff.

### 2.6 Network interpretation

We adopted a widely-used visualization method [7, 24] to convert the first layer kernels to PWMs. For each convolutional kernel, we searched through all the samples for all that can activate at least one neuron (output of the neuron > 0.5 of the maximum output among all samples) in the first convolutional layer. Each such activation was mapped back to the input sequence to locate the region that led to the activation. For each convolutional kernel, we aligned all of the activating sequences to generate a PWM. To understand the biological meaning of these PWMs, we used tomtom ([25], ver 4.11.1) to match the PWMs to known human motifs in CIS-BP database [26] with a FDR threshold of 0.1 as suggested in Kelly et al. [7] When combined with importance analysis, we used a more stringent FDR of 0.01. For the analysis of PWMS partially matched with known motifs, we also compared the PWMs against TransFac [27] database.

We interpret the importance of the first layer kernels with an optimization-based framework. We fixed all weights in a trained CpGenie model, and optimized the output of the neuron in the last layer that corresponds to the target label (methylated/unmethylated, or tissue-specific/tissue-invariant) with respect to the input of the second layer (i.e. the output of the first max-pooling layer) under a *L*_2_ regularization. The resulting optimum input is a 2*D* matrix, representing the spatial activation pattern of each of the first layer convolutional kernels for the network to reach high confidence in the prediction. For each kernel, we assigned the importance as the maximum activation from all locations.

## 3 Results

### 3.1 A two-step variant-evaluation framework for DNA methylation

CpGenie employs a two-step framework to evaluate the impact of genetic variants on DNA methylation. The first module (Me-CpG-prediction) predicts the DNA methylation status of a CpG from its flanking 1001bp sequence context. As a fully sequence-based model, Me-CpG-prediction learns a regulatory code of DNA methylation from genomic sequence, which is essential for accurate allele-specific predictions and non-coding variant evaluation. The second module (variant-prediction) uses the regulatory code learned in Me-CpG-prediction to score the impact of a given genetic variant on proximal region. Variant-prediction predicts the methylation modulation caused by a variant by summarizing diverse statistics of the predicted methylation changes in adjacent CpG sites.

Me-CpG-prediction employs a convolutional neural network (CNN) to learn the sequence determinants for DNA methylation. Compared to random forest or support-vector machines (SVM) methods which are often used in existing frameworks [17, 19, 20, 28, 29], a CNN is able to learn more effectively from large-scale DNA methylation datasets, such as WGBS and RRBS, and is capable to learn features of different spatial and complexity scales using the hierarchical architecture.

Variant-prediction applies Me-CpG-prediction to characterize the impact of a genetic variant across a region. Variant-prediction first uses Me-CpG-prediction to score the impact of a variant compared with the corresponding wild-type allele on all CpGs within 500bp of the variant. Variant-prediction then scores a variant’s impact by the change in the sum, max and mean of methylation level in a 1001bp genomic neighborhood around the variant when compared with the wild-type allele. These different statistics, which are often not correlated, describe different aspects of the impact and in sum produce a succinct yet informative picture of how a sequence variant alters the local methylation landscape.

### 3.2 CpGenie predicts DNA methylation from sequence context

We first assessed the ability of CpGenie’s Me-CpG-prediction module to predict the methylation status of a CpG site from its flanking sequence. As none of the published models for fully sequence-based DNA methylation prediction [17–20] provide an standalone software for re-training and predicting on a large number of CpG sites, we compared CpGenie with a random forest (RF) classifier trained on 4-mer frequencies of the input sequence, considering that random forest and k-mer frequencies have been used in the literature of sequence-based DNA methylation models [19, 20, 28, 29].

We evaluated Me-CpG-prediction and the random forest method on restricted representation bisulfite sequencing (RRBS) datasets from ENCODE. We trained Me-CpG-prediction and the random forest method on RRBS data from GM12878, a lymphoblastoid cell line (LCL) extensively studied in ENCODE. Me-CpG-prediction resulted in an area under receiver operating characteristic (auROC) of 0.854 and an area under precision-recall curve (auPRC) of 0.685 on the held-out test set. Both of these metrics surpassed the performance of the random forest baseline (auROC of 0.814 and auPRC of 0.584, Figure 2A).

We then evaluated Me-CpG-prediction on 50 RRBS datasets from ENCODE (Supplementary Table 1) with our random forest baseline to systematically benchmark their capacity in predicting DNA methylation. Me-CpG-prediction robustly outperformed the alternative methods, with better auROC and auPRC for all 50 experiments (Figure 2C). We also found that a sequence window of 1001bp optimized performance (Supplementary Figure 1), which could suggest stronger involvement of sequence features within 500bp away in DNA methylation regulation.

We further characterized Me-CpG-prediction and competing methods using a bisulfite sequencing dataset from LCLs derived from 60 Yoruban (YRI) HapMap individuals [21]. Me-CpG-prediction achieved an auROC of 0.75 and an auPRC of 0.79 (Figure 2B), surpassing the competing model trained and tested on the same datasets (auROC = 0.57, auPRC = 0.63).

### 3.3 CpGenie predicts the impact of functional variants on DNA methylation

We next assessed the ability of CpGenie’s variant-prediction module to identify genetic variants that modulate DNA methylation. Kaplow et al. analyzed the DNA methylation level of over 800,000 single nucleotide polymorphism (SNP)-CpG pairs by mapping the bisulfite sequencing reads back to the reference and alternate allele of a variant [21]. They found over 2,000 genetic variants (meQTLs) with statistically significant allelic imbalance of DNA methylation. As only reads that overlap with both the CpG site and the variant locus were counted, the meQTLs discovered from this method act *in cis* (with an average distance of 25.4 bp), making these data a relevant standard to evaluate the ability to predict allelic change of DNA methylation in the presence of a sequence variant.

Variant-prediction accurately predicted the direction of allelic methylation change caused by sequence variants. When applied to the meQTLs on chromosomes held out from training, variant-prediction accurately identified the allele with higher DNA methylation level, showing sensitivity and accuracy to single-nucleotide changes (Figure 3A). Moreover, the accuracy quickly and stably increased to 100% when we gradually retained only the high-confidence predictions by increasing the threshold of the absolute allelic difference in the predicted methylation (Figure 3B). For instance, for the variants of which the predicted absolute difference of DNA methylation between the two alleles is greater than 0.03, CpGenie identified the allele with more methylation with an accuracy > 90%.

We find that CpGenie variant-prediction accurately classifies variants that are meQTLs from variants that exhibit no impact on DNA methylation. Since CpGenie is the first computational method to predict meQTLs we compare it with several state-of-the-art methods for functional variant prioritization, including DeepSEA [6], Basset [7], deltaSVM [8], GWAVA [5], and CADD [4]. We used 201 meQTLs on the chromosomes held out in the training of CpGenie models as positive samples. To simulate different equilibrium linkage structures we constructed three negative sample sets that are 10 times, 50 times, and 100 times the size of the positive set by randomly sampling from the 76,532 non-meQTLs on the held-out chromosomes. CpGenie models trained on datasets from Kaplow et al and ENCODE GM12878 RRBS datasets both surpassed the competing methods, with higher accuracy at 10% recall and larger area under precision recall curve (auPRC) when evaluated on all three negative sets (Figure 3C). Thus, CpGenie excels in predicting genetic variants that module DNA methylation, an important task that the state-of-the-art frameworks for functional variant prioritization fail in.

### 3.4 CpGenie learns the binding motifs of proteins known to regulate DNA methylation

We expected that a predictive model of DNA methylation from sequence would learn motifs that correspond to regulators associated with the mechanism of DNA methylation. The basic unit of a convolutional layer is a “kernel” that searches for patterns in the input, analogous to a motif scanner looking for motif matches. Interpreting the convolutional kernels in the first layer of a network is crucial for understanding how the network responds to an input sequence [7, 24]. Previous studies have established that many transcription factors interact with DNA methyltransferases (DNMT) that methylate DNA [30]. As transcription factors are known for binding DNA with strong sequence specificity, we transformed the first-layer convolutional kernels in CpGenie to position weight matrices (PWMs) (Methods) to determine if it learned certain of these sequence motifs.

We found that 97 out of the 128 PWMs recognized by CpGenie’s Me-CpG-prediction significantly match the motifs of known transcription factors (Figure 4A, Supplemental Table 2). We found the motifs of 21 transcription factors known to strongly interact with a DNA methyltransferase (DMNT) [30], including ELK1,FLI1 and E2F4. As a point of comparison, Das et al. [31] found that the motifs of 31 transcription factors, only a small fraction (6/31) of which overlap with the DNMT-interacting factors, help classify hyper-methylated regions from hypo-methylated regions. We found 48% (15/31) of the Das et al. discovered motifs were among the CpGenie-discovered PWMs including YY1 and CEBPA. Moreover, although without a statistically significant overall match, many CpGenie discovered PWMs capture motif information associated with transcription factors previously reported to be associated with DNA methylation, such as NFKB1, MEF3, and LUN1 (Figure 4B).

Interestingly, a large number of CpGenie’s Me-CpG-prediction discovered PWMs are variants of PAX4 and SP3 motifs (24 and 23 respectively). Hervouet et al. reported DNMT-interaction with other transcription factors in the same family (PAX6, SP1 and SP4). Certain of the predictive transcription factor motifs discovered with CpGenie are not known for involvement in DNA methylation. Two examples are GFI1 (FDR q-value=0.0025), which is a transcriptional repressor that functions by histone deacetylase (HDAC) recruitment, and THRA (FDR q-value=0.0023), which is a nuclear hormone receptor that can act as a repressor or activator of transcription.

We next scored the importance of CpGenie’s Me-CpG-prediction 128 first-layer convolution kernels with an optimization-based framework (Methods). The framework identifies the first-layer kernel activation pattern that can maximize the network’s confidence to classify a sample as one class (methylated / unmethylated). To understand the biological relevance of the top-ranking kernels, we chose a more stringent false discovery rate of 0.01 when matching with known motifs. The top 10 convolution kernels for high and low methylation prediction are quite distinct, with the exception that SP3 is important for predicting both high and low methylation (Supplementary Table 3).

### 3.5 CpGenie assists in downstream analysis of functional variants

We next asked whether CpGenie, which is optimized for meQTL prediction, could shed light on the functional consequence of genetic variants associated with downstream phenotypes. We applied CpGenie on four experimentally validated GWAS SNPs, rs1427407 (fetal hemoglobin levels, [32]), rs12740374 (low-density lipoprotein (LDL) cholesterol levels,[33]), rs10737680 (age-related macular degeneration, [34]), rs7705033 (Visceral adipose tissue, [35]), of which the first two have been reported to alter gene expression (BCL11A [36] and SORT1 [37] respectively) and the last two have been reported to alter DNA methylation [21]. Compared with linked SNPs in strong linkage disequilibrium (Methods), both the validated SNPs were scored higher by CpGenie (Figure 5A).

We further applied CpGenie on two much larger GWAS SNPs and eQTL datasets [6], one with 78,613 eQTLs from GRASP (Genome-Wide Repository of Associations between SNPs and Phenotypes) [38] and one with 12,296 disease-associated SNPs from the US National Human Genome Research Institute’s GWAS Catalog [39]. For each dataset, five size-matched negative sets were constructed by sampling from different subsets of 1000 Genome Project SNPs [40]. We found a simple L2-regularized logistic regression model trained on CpGenie’s predictions for the 50 ENCODE RRBS datasets performed competitively in both eQTL and GWAS SNP prioritization (Figure 5B), compared to several state-of-the-art methods including DeepSEA [6], CADD [4], GWAVA [5] and Funseq2 [41] which were all trained on more diverse sets of functional data such as histone modification, transcription factor binding, and gene expression.

To assess the relative importance of DNA methylation features in eQTL and GWAS SNPs prediction, we combined the functional features predicted from CpGenie and DeepSEA, and trained a random forest model in which features importance can be evaluated by mean decrease impurity (Methods). DeepSEA predicts a variant’s effect by producing 919 features derived from experiments of DNase hyper-sensitivity (DNase-seq), transcription factor binding (ChIP-seq), and histone modification (ChIP-seq), which is much larger and comprehensive than CpGenie’s prediction of methylation alone. However, in both eQTL and GWAS SNP prioritization, CpGenie-predicted DNA methylation features are considered significantly more important than the original DeepSEA features as a whole (Mann-Whitney U test, Figure 5C, Supplemental Table 4). To account for the potential inflation of significance from the high correlation in the features, we further only looked at the top features with significantly higher importance (Methods). Consistently across different tasks, CpGenie-predicted methylation-based features account for a significant portion in the top features which the eQTL/GWAS predictor might actually rely on (Supplemental Table 4).

## 4 Discussion

Despite the growing number of genetic variants associated with disease and complex traits by genome-wide association studies (GWAS), the identification of causal variants and their pathogenic mechanisms remains a challenge that requires predictive models for accurate interpretation of non-coding variants. CpGenie is a computational framework that is able to assess non-coding variant’s effect on DNA methylation, a functional signal largely overlooked by existing models for functional variant prioritization.

With its convolutional neural network-based Me-CpG-prediction, CpGenie is able to learn sophisticated sequence determinants associated with DNA methylation efficiently from large-scale DNA methylation data generated from high-throughput bisulfite sequencing technology. CpGenie predicts the DNA methylation status of a CpG solely from the sequence context with consistently high accuracy on datasets across different cell lines, tissues, and experiment protocols (Figure 2), demonstrating crucial robustness and generalizability of the methodology.

CpGenie demonstrated high sensitivity to single-base changes in input sequences (Figure 3AB), a capability that enables the incorporation of allele-specific, rather than regional, DNA methylation information in the interpretation of non-coding variants. CpGenie identifies methylation quantitative trait loci (meQTLs) with a precision that surpasses the state-of-the-art frameworks for functional variants prioritization (Figure 3C), which emphasizes the unique role of CpGenie in providing a more comprehensive and accurate interpretation of the functional consequence of non-coding variants.

We have shown that CpGenie’s methylation-based predictions can assist in the downstream analysis of risk-associated variants. By taking into account the influence on all the CpG sites residing in the neighborhood of the target variant, CpGenie summarizes a diverse set of statistics that can both directly instruct the identification of causal variant from candidates in strong linkage disequilibrium (Figure 5A) and be incorporated in secondary models designed to classify sequence variants associated with gene expression or complex traits (Figure 5B). Moreover, CpGenie’s methylation-based features are highly favored (Figure 5C), when jointly considered with other functional annotations, such as DNase hyper-sensitivity, histone marks and transcription factor binding, by a predictive model for functional variant classification. This demonstrates the wealth of information captured in the change of DNA methylation and highlights the necessity to include allele-specific DNA methylation predictions in a comprehensive assessment of non-coding variants.

We envision CpGenie to be a resource to help understand the regulatory mechanism encoded in the non-coding region of the genome, and contribute to the functional interpretation of non-coding variants associated with complex traits and diseases.

## Acknowledgments

We acknowledge helpful input from Professor Brendan J. Frey and his lab. We are grateful for insights and suggestions from other members in Gifford Lab.

---

Figure 1: Schematics of CpGenie. (A) CpGenie takes the high-throughput DNA methylation sequencing data, such as restricted representation bisulfite sequencing (RRBS) or whole-genome bisulfite sequencing (WGBS) as input and produces predictions of CpG methylation as output. CpGenie can predict DNA methylation at CpG resolution, interpreting the functional consequence of non-coding sequence variants, and prioritizing causal mutations from GWAS-determined associations. (B) CpGenie converts the sequence context around a CpG into one-hot encoding, and transforms it to higher-level features through three pairs of convolutional and max-pooling layers. Two fully-connected layers follow to make predictions on the methylation status of the queried CpG.

Figure 2: CpGenie predicts DNA methylation at CpG resolution. (A,B) The receiver operating characteristic (ROC) curve (top) and precision-recall (PRC) curve (bottom) of CpGenie (blue) and random forest using 4-mer counts (green) for predicting DNA methylation status of held-out CpGs in GM12878 RRBS data (A) and bisulfite sequencing data from LCLs derived from 60 Yoruban HapMap individuals (B). (C) Pairwise auROC (top) and auPRC (bottom) comparison of CpGenie (y-axis) and random forest using 4-mer counts (x-axis) on 50 RRBS datasets from ENCODE.

Figure 3: CpGenie accurately predicts the direction of allele-specific (AS) DNA methylation and prioritizes variants that module DNA methylation (meQTLs). (A) CpGenie’s DNA methylation prediction for the reference and alternate alleles of 201 meQTLs on held-out chromosome 11 and 12. The x and y axis represents the CpGenie predicted DNA methylation level. The blue and red dots represent reference allele-biased and alternate allele-biased variants respectively as experimentally determined by Kaplow et al. (B) Prediction accuracy quickly and steadily increased to 100% when only the high-confidence predictions were retained. The y-axis denotes accuracy and the x-axis represents margin, or the threshold of predicted absolute allelic difference in methylation to retain high-confidence predictions. (C) The precision-recall curve (PRC) for classifying the 201 meQTL from three different random subsets of the 76,532 non-meQTL that are 10 times (left), 50 times (middle), and 100 times (right) the size of meQTL. CpGenie outperformed all the state-of-the-art methods in functional variant prioritization with better precision at the 10% recall and higher area under precision-recall curve.

Figure 4: CpGenie learns motifs of regulatory elements involved in DNA methylation. (A) 97 out of 128 of the convolutional filters match motifs of known transcription factors in the human CIS-BP database at an FDR threshold of 0.1.(B) Examples of convolutional kernels characterizing partial information of transcription factors known for involvement in or predictive for DNA methylation. The logos for LUN1 and MEF3 were generated from motif information in TransFac databse (Jan. 2013) and the logo for NFKB1 was generated from motif information in CIS-BP database.

Figure 5: CpGenie’s sequence-based DNA methylation predictions assist in downstream analysis of functional variants. (A) CpGenie score the validated GWAS SNPs (red) higher than the SNPs in strong linkage disequilibrium. The three statistics generated from CpGenie are colored in blue (the absolute change of total methylation of proximal CpG sites), green (the absolute change of mean methylation of proximal CpG sites) and red (the absolute change of maximum methylation of proximal CpG sites). (B) Compared to previous methods that utilize more annotation information, CpGenie achieved better or comparable performance in prioritizing noncoding GRASP eQTLs (left) and noncoding GWAS Catalog SNPs (right) against noncoding 1000 Genome Project SNPs. The x-axis denotes the mean distance of the SNPs in the negative set to the paired positive SNP. The ‘Random’ group denotes 1,000,000 randomly sampled 1000 Genome Project SNPs. (C) CpGenie’s DNA methylation features (green) were considered significantly more important in general than DeepSEA’s functional predictions on histone modification, transcription factor binding and DNase hypersensitivity (blue) in eQTL (left) and GWAS SNPs (right) prioritization. The asterisks denote statistical significance calculated from Mann-Whitney U test (p-value < 0.001).

## References

[1] Lucia a Hindorff, Praveen Sethupathy, Heather a Junkins, Erin M Ramos, Jayashri P Mehta, Francis S Collins, and Teri a Manolio. Potential etiologic and functional implications of genome-wide association loci for human diseases and traits. Proceedings of the National Academy of Sciences of the United States of America, 106(23):9362–7, June 2009.

[2] Matthew T Maurano, Richard Humbert, Eric Rynes, Robert E Thurman, Eric Haugen, Hao Wang, Alex P Reynolds, Richard Sandstrom, Hongzhu Qu, Jennifer Brody, et al. Systematic localization of common disease-associated variation in regulatory dna. Science, 337(6099):1190–1195, 2012.

[3] Alexander Gusev, S Hong Lee, Gosia Trynka, Hilary Finucane, Bjarni J Vilhjálmsson, Han Xu, Chongzhi Zang, Stephan Ripke, Brendan Bulik-Sullivan, Eli Stahl, et al. Partitioning heritability of regulatory and cell-type-specific variants across 11 common diseases. The American Journal of Human Genetics, 95(5):535–552, 2014.

[4] Martin Kircher, Daniela M Witten, Preti Jain, Brian J O’Roak, Gregory M Cooper, and Jay Shendure. A general framework for estimating the relative pathogenicity of human genetic variants. Nature genetics, 46(3):310, 2014.

[5] Graham RS Ritchie, Ian Dunham, Eleftheria Zeggini, and Paul Flicek. Functional annotation of noncoding sequence variants. Nature methods, 11(3):294–296, 2014.

[6] Jian Zhou and Olga G Troyanskaya. Predicting effects of noncoding variants with deep learning-based sequence model. Nature methods, 12(10):931–934, aug 2015.

[7] David R Kelley, Jasper Snoek, and John L Rinn. Basset: Learning the regulatory code of the accessible genome with deep convolutional neural networks. Genome research, 2016.

[8] Dongwon Lee, David U Gorkin, Maggie Baker, Benjamin J Strober, Alessandro L Asoni, Andrew S McCallion, and Michael a Beer. A method to predict the impact of regulatory variants from DNA sequence. Nature Genetics, (June), 2015.

[9] Adrian Bird. Dna methylation patterns and epigenetic memory. Genes & development, 16(1):6–21, 2002.

[10] Christoph Bock. Analysing and interpreting dna methylation data. Nature Reviews Genetics, 13(10):705–719, 2012.

[11] Denise P Barlow. Genomic imprinting: a mammalian epigenetic discovery model. Annual review of genetics, 45:379–403, 2011.

[12] Marion Martin and Zdenko Herceg. From hepatitis to hepatocellular carcinoma: a proposed model for cross-talk between inflammation and epigenetic mechanisms. Genome medicine, 4(1):1, 2012.

[13] Alexander Meissner. Epigenetic modifications in pluripotent and differentiated cells. Nature biotechnology, 28(10):1079–1088, 2010.

[14] Timothy H Bestor. The host defence function of genomic methylation patterns. In Novartis Found. Symp, volume 214, pages 187–195, 1998.

[15] Hyung Joo Lee, Rebecca F Lowdon, Brett Maricque, Bo Zhang, Michael Stevens, Daofeng Li, Stephen L Johnson, and Ting Wang. Developmental enhancers revealed by extensive dna methylome maps of zebrafish early embryos. Nature communications, 6, 2015.

[16] Woochang Hwang, Verity F Oliver, Shannath L Merbs, Heng Zhu, and Jiang Qian. Prediction of promoters and enhancers using multiple dna methylation-associated features. BMC genomics, 16(7):1, 2015.

[17] Manoj Bhasin, Hong Zhang, Ellis L Reinherz, and Pedro A Reche. Prediction of methylated CpGs in DNA sequences using a support vector machine. FEBS letters, 579(20):4302–8, aug 2005.

[18] S Kim, M Li, H Paik, K Nephew, H Shi, R Kramer, D Xu, and T H Huang. Predicting DNA methylation susceptibility using CpG flanking sequences. In Pacific Symposium on Biocomputing. Pacific Symposium on Biocomputing, pages 315–26, jan 2008.

[19] Lingyi Lu, Kao Lin, Ziliang Qian, Haipeng Li, Yudong Cai, and Yixue Li. Predicting DNA methylation status using word composition. Journal of Biomedical Science and Engineering, 03(07):672–676, jul 2010.

[20] Xuan Zhou, Zhanchao Li, Zong Dai, and Xiaoyong Zou. Prediction of methylation CpGs and their methylation degrees in human DNA sequences. Computers in biology and medicine, 42(4):408–13, apr 2012.

[21] Irene M Kaplow, Julia L MacIsaac, Sarah M Mah, Lisa M McEwen, Michael S Kobor, and Hunter B Fraser. A pooling-based approach to mapping genetic variants associated with dna methylation. Genome research, 25(6):907–917, 2015.

[22] Guillaume Marçais and Carl Kingsford. A fast, lock-free approach for efficient parallel counting of occurrences of k-mers. Bioinformatics, 27(6):764–770, 2011.

[23] Leo Breiman, Jerome Friedman, Charles J Stone, and Richard A Olshen. Classification and regression trees. CRC press, 1984.

[24] Babak Alipanahi, Andrew Delong, Matthew T Weirauch, and Brendan J Frey. Predicting the sequence specificities of DNA - and RNA-binding proteins by deep learning. Nature biotechnology, 33(8):831–838, jul 2015.

[25] Shobhit Gupta, John A Stamatoyannopoulos, Timothy L Bailey, and William Stafford Noble. Quantifying similarity between motifs. Genome biology, 8(2):1, 2007.

[26] Matthew T Weirauch, Ally Yang, Mihai Albu, Atina G Cote, Alejandro Montenegro-Montero, Philipp Drewe, Hamed S Najafabadi, Samuel A Lambert, Ishminder Mann, Kate Cook, et al. Determination and inference of eukaryotic transcription factor sequence specificity. Cell, 158(6):1431–1443, 2014.

[27] Vea Matys, Ellen Fricke, R Geffers, Ellen Gößling, Martin Haubrock, R Hehl, Klaus Hornischer, Dagmar Karas, Alexander E Kel, Olga V Kel-Margoulis, et al. Transfac®: transcriptional regulation, from patterns to profiles. Nucleic acids research, 31(1):374–378, 2003.

[28] Weiwei Zhang, Tim D Spector, Panos Deloukas, Jordana T Bell, and Barbara E Engelhardt. Predicting genome-wide DNA methylation using methylation marks, genomic position, and DNA regulatory elements. Genome biology, 16(1):14, jan 2015.

[29] Shicai Fan, Kang Huang, Rizi Ai, Mengchi Wang, and Wei Wang. Predicting CpG methylation levels by integrating Infinium HumanMethylation450 BeadChip array data. Genomics, 2016.

[30] Eric Hervouet, François M Vallette, and Pierre-François Cartron. Dnmt3/transcription factor interactions as crucial players in targeted dna methylation. Epigenetics, 4(7):487–499, 2009.

[31] Rajdeep Das, Nevenka Dimitrova, Zhenyu Xuan, Robert A Rollins, Fatemah Haghighi, John R Edwards, Jingyue Ju, Timothy H Bestor, and Michael Q Zhang. Computational prediction of methylation status in human genomic sequences. Proceedings of the National Academy of Sciences of the United States of America, 103(28):10713–6, jul 2006.

[32] Siana Nkya Mtatiro, Tarjinder Singh, Helen Rooks, Josephine Mgaya, Harvest Mariki, Deogratius Soka, Bruno Mmbando, Evarist Msaki, Iris Kolder, Swee Lay Thein, et al. Genome wide association study of fetal hemoglobin in sickle cell anemia in tanzania. PloS one, 9(11):e111464, 2014.

[33] Sekar Kathiresan, Cristen J Willer, Gina M Peloso, Serkalem Demissie, Kiran Musunuru, Eric E Schadt, Lee Kaplan, Derrick Bennett, Yun Li, Toshiko Tanaka, et al. Common variants at 30 loci contribute to polygenic dyslipidemia. Nature genetics, 41(1):56–65, 2009.

[34] AMD Gene Consortium et al. Seven new loci associated with age-related macular degeneration. Nature genetics, 45(4):433–439, 2013.

[35] Caroline S Fox, Yongmei Liu, Charles C White, Mary Feitosa, Albert V Smith, Nancy Heard-Costa, Kurt Lohman, Andrew D Johnson, Meredith C Foster, Danielle M Greenawalt, et al. Genome-wide association for abdominal subcutaneous and visceral adipose reveals a novel locus for visceral fat in women. PLoS Genet, 8(5):e1002695, 2012.

[36] Daniel E Bauer, Sophia C Kamran, Samuel Lessard, Jian Xu, Yuko Fujiwara, Carrie Lin, Zhen Shao, Matthew C Canver, Elenoe C Smith, Luca Pinello, et al. An erythroid enhancer of bcl11a subject to genetic variation determines fetal hemoglobin level. Science, 342(6155):253–257, 2013.

[37] Kiran Musunuru, Alanna Strong, Maria Frank-Kamenetsky, Noemi E Lee, Tim Ahfeldt, Katherine V Sachs, Xiaoyu Li, Hui Li, Nicolas Kuperwasser, Vera M Ruda, et al. From noncoding variant to phenotype via sort1 at the 1p13 cholesterol locus. Nature, 466(7307):714–719, 2010.

[38] Richard Leslie, Christopher J O’Donnell, and Andrew D Johnson. Grasp: analysis of genotype-phenotype results from 1390 genome-wide association studies and corresponding open access database. Bioinformatics, 30(12):i185–i194, 2014.

[39] Danielle Welter, Jacqueline MacArthur, Joannella Morales, Tony Burdett, Peggy Hall, Heather Junkins, Alan Klemm, Paul Flicek, Teri Manolio, Lucia Hindorff, et al. The nhgri gwas catalog, a curated resource of snp-trait associations. Nucleic acids research, 42(D1):D1001–D1006, 2014.

[40] 1000 Genomes Project Consortium et al. An integrated map of genetic variation from 1,092 human genomes. Nature, 491(7422):56–65, 2012.

[41] Yao Fu, Zhu Liu, Shaoke Lou, Jason Bedford, Xinmeng Jasmine Mu, Kevin Y Yip, Ekta Khurana, and Mark Gerstein. Funseq2: a framework for prioritizing noncoding regulatory variants in cancer. Genome biology, 15(10):1, 2014.

